# Antifungal siderophore conjugates for theranostic applications in invasive pulmonary aspergillosis using low molecular TAFC scaffolds

**DOI:** 10.1101/2021.04.19.440472

**Authors:** Joachim Pfister, Milos Petrik, Katerina Bendova, Barbara Matuszczak, Ulrike Binder, Alexander Kühbacher, Fabio Gsaller, Matthias Misslinger, Hubertus Haas, Clemens Decristoforo

## Abstract

Invasive pulmonary aspergillosis (IPA) is a life-threatening form of fungal infection, primarily in immunocompromised patients and associated with a significant mortality. Diagnostic procedures are often invasive and/or time consuming and existing antifungals can be constrained by dose limiting toxicity and drug interaction. In this study, we modified triacetylfusarinine C (TAFC), the main siderophore produced by the opportunistic pathogen *Aspergillus fumigatus*, with antifungal molecules to perform antifungal susceptibility tests and molecular imaging.

**Methods:** A variation of small organic molecules (eflornithine, fludioxonil, thiomersal, fluoroorotic acid (FOA), cyanine 5 (Cy5)) with antifungal activity were coupled to TAFC, resulting in a “Trojan horse” to deliver antifungal compounds specifically into *Aspergillus fumigatus* hyphae by the major facilitator transporter MirB. Radioactive labelling with gallium-68 allowed to perform *in vitro* characterization (LogD, stability, uptake assay) as well as biodistribution experiments and PET/CT imaging in an IPA rat infection model. Compounds labelled with stable gallium were used for antifungal susceptibility tests.

**Results:** [Ga]DAFC-fludioxonil, -FOA and Cy5 revealed a MirB dependent active uptake with fungal growth inhibition at 16 μg/mL after 24 h. Visualization of an *Aspergillus fumigatus* infection in lungs of a rat was possible with gallium-68 labelled compounds using PET/CT. Heterogeneous biodistribution patterns revealed the immense influence of the antifungal moiety conjugated to DAFC.

**Conclusion:** Overall, novel antifungal siderophore conjugates with promising fungal growth inhibition and the possibility to perform PET-imaging, combine both therapeutic and diagnostic potential in a theranostic compound for IPA caused by *Aspergillus fumigatus*.

## 1. Introduction

Invasive aspergillosis (IA) is a severe infection in humans, associated with high mortality and an estimated incidence of 250,000 cases a year worldwide [1,2]. IA most commonly involves the lungs resulting in invasive pulmonary aspergillosis (IPA) in immunocompromised patients with *Aspergillus fumigatus* (*A. fumigatus*) as the most common opportunistic pathogen involved [3,4].

Diagnostic procedures recommended by current guidelines [5] are often invasive (e.g. bronchoalveolar lavage, bronchoscopy) and/or time consuming, worsen survival outcome for patients [2,6]. Current treatment options are azoles, echinocandins and polyenes [5,7] but antifungal resistance is a concern for the management of *A. fumigatus* infections [8]. Furthermore, antifungal agents are often constrained by drug interaction, route of administration and dose-limiting toxicity. These factors are very alarming and demand for a continuous improvement in the diagnosis and treatment of IPA.

During infection, the human body sequesters iron as a defense strategy to prevent growth of pathogens [9]. Iron is an essential micronutrient and *A. fumigatus* has developed sophisticated strategies to overcome this problem: low-affinity iron uptake, reductive iron assimilation (RIA) and siderophore-mediated iron acquisition (SIA) [10] whereby RIA and SIA are high-affinity iron uptake systems. SIA is highly upregulated during infection and essential for the virulence of *A. fumigatus* [11]. Siderophores are low-molecular mass organic molecules with a high affinity to bind ferric iron (Fe(III)). *A. fumigatus* produces different types of siderophores: desferry-fusarinine C (FsC) and desferry-triacetylfusarinine C (TAFC) for external iron acquisition and desferri-ferricrocin, desferri-hydroxy-ferricrocin for internal iron storage [12]. After binding iron in the environment, [Fe]TAFC is taken up by the hyphae with a specific transporter called MirB [13]. This transporter is highly upregulated during infection [11] and additionally, [Fe]TAFC shows a very high specificity for *A. fumigatus* lacking uptake by mammalian cells [14,15]. This offers an opportunity to use MirB as a target for imaging and treatment of *A. fumigatus* infections.

Petrik et al. showed the possibility of exchanging iron from [Fe]TAFC with the radioactive isotope gallium-68 to perform PET/CT imaging in an lung infection animal model [16]. Furthermore, by using diacetylfusarinine C ([Fe]DAFC), which possesses a free amine group, chemical modifications are possible [17]. In previous experiments by our group, we used fluorescent dyes coupled to DAFC to perform fluorescence-microscopy of *A. fumigatus* hyphae [18] and labelled with gallium-68, hybrid imaging of an infection in a rat model [19]. This proof of principle using TAFC as a “Trojan horse” to channel a variety of molecules specifically into *A. fumigatus* via the MirB transporter, led us to the idea to adopt this system for antifungal molecules. For this purpose, different candidates (Figure 1) were chosen depending on certain, individual properties.

**Figure 1.**
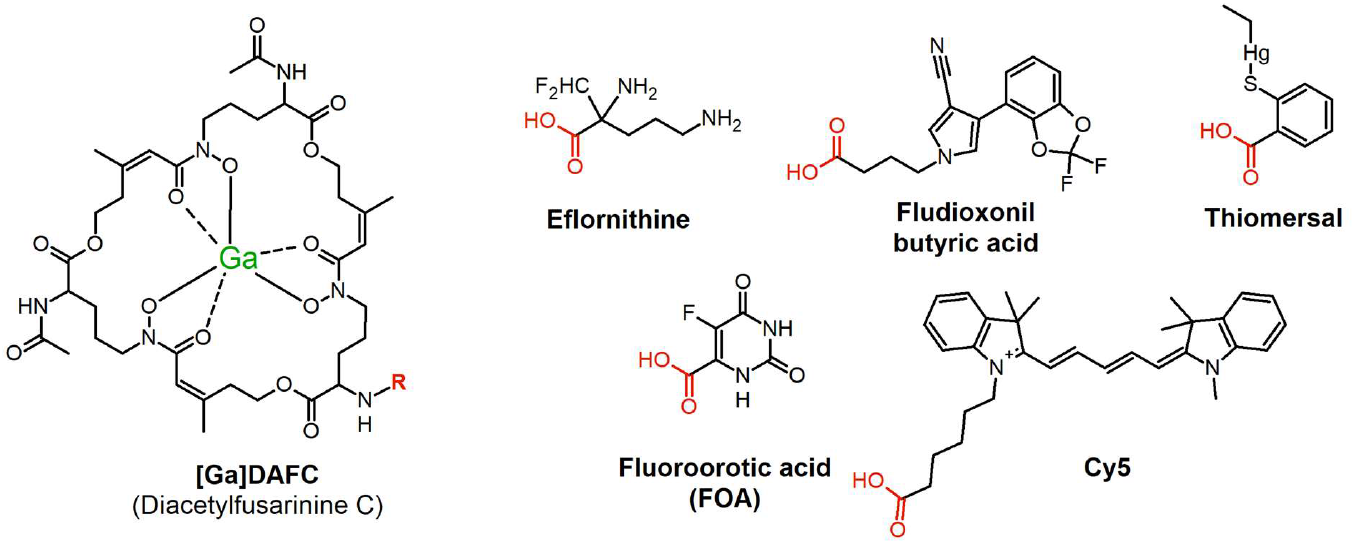
Chemical structures of antifungal compounds used in this study. On the left side: gallium labelled form of diacetylfusarinine C (DAFC); red part of the structures indicate coupling site of individual molecules.

Eflornithine, an inhibitor of ornithine decarboxylase [20], is a very multifunctional drug as it is used topically (15% eflornithine cream) against unwanted facial hair (hirsutism) [21], as a treatment for malignant gliomas [22] and as intravenous cocktail in combination with nifurtimox, approved by the EMA against African trypanosomiasis (sleeping sickness) caused by *Trypanosoma brucei gambiense* [23,24]. Beckmann et al. showed an antifungal activity against *A. fumigatus* in agar diffusion assays [25], which potentially could be enhanced by active transport into the hyphae via MirB. The same rational was applied for fluoroorotic acid, which is converted by ornithine-5’-monophosphate (OMP)-decarboxylase to the toxic intermediate 5-fluoro-UMP, commonly used for selection in yeast genetic experiments [26]. Furthermore, we chose fludioxonil, a phenylpyrrole pesticide used post-harvest for fruit and vegetable crops to minimize losses from mold contamination during transport and point of sale. It inhibits class III hybrid histidine kinases (HHK) that are peculiar to fungi and regulate the high osmolarity glycerol pathway (HOG) [27]. For the last 30 years it has been postulated to be safe for human as there is no target in the body, but studies are currently ongoing to investigate if there are adverse health effects on the cellular level [28]. By coupling fludioxonil to DAFC, the active uptake by MirB could lower the administered dosage needed and interaction with human targets could be hindered by this modification. A similar concept could be applied for thiomersal which was commonly used as a preservative substance in vaccines and ophthalmic products [29,30] but adverse health effects of ethylmercury (active metabolite) are still controversially discussed [31]. The last compound Cy5 was originally designed as an agent for hybrid imaging and microscopy, whereby an antifungal activity was observed during growth experiments by our group [18,19].

The aim of the study was to investigate the growth inhibition potential of various antifungal siderophore conjugates and to visualize an *A. fumigatus* lung infection via PET/CT in a rat model by labelling the compounds with gallium-68. Thereby, beside antifungal properties also pharmacokinetic differences of individual compounds were revealed, providing essential data for further applications aiming at MirB as a drug target. This proof of principle shows the possibility of using these compounds for therapy and diagnostic with the same molecule, in a so called theranostic approach.

## 2. Materials and Methods

### 2.1. Chemicals and Synthesis of Antifungal conjugates

All chemicals were purchased from Sigma Aldrich as reagent grade and used without further purification unless stated otherwise. Exact synthetic procedures and analytical details can be found in the supplementary material. Starting from iron containing diacetylfusarinine C ([Fe]DAFC) all antifungal conjugates were conjugated forming an amide bond using O-(7-Azabenzotriazol-1-yl)-N,N,N’,N’-tetramethyluronium hexafluorophosphate (HATU) as an activating reagent. To couple eflornithine, free amino groups in the molecular structure had to be protected with tert-butyloxycarbonyl (Boc) before linking with [Fe]DAFC followed by cleavage of the Boc protecting groups with trifluoroacetic acid. fludioxonil had to be modified with ethyl 4-bromobutyrate resulting in a new compound with an ester function, followed by hydrolysis to get the free free carboxylic acid, suitable for HATU coupling.

All siderophore conjugates were purified by preparative HPLC and freeze dried. For iron removal, compounds were incubated with 100 mM Na_2_EDTA solution and subsequently purified by preparative HPLC, resulting in the iron free form for radiolabelling experiments. Stable gallium containing siderophore conjugates were produced by coupling the antifungal directly to [Ga]DAFC to reduce iron contamination to a minimum. Exact coupling conditions and a detailed description can be found in the supplementary material as well as analytical data for each compound, respectively.

#### Radiolabelling

Gallium-68 was produced by fractionated elution of ^68^Ge/^68^Ga-generator (IGG100. Eckert & Ziegler Isotope Products, Berlin, Germany; nominal activity of 1850 MBq) with 0.1 M hydrochloric acid (HCL, Rotem Industries, Arva, Israel). For labelling, 10 μg (5–8 nmol) of DAFC-conjugate were mixed with 200 μL gallium eluate (~15–30 MBq) and the pH was adjusted to 4.5 by adding 20 μL of sodium acetate solution (1.14 M) per 100 μL eluate. The mixture was left to react for 10 min at RT and finally analyzed by radio-TLC and radio-RP-HPLC [32].

##### *In Vitro* Experiments

###### Distribution Coefficient (LogD)

To determine the lipophilicity of the conjugates, the distribution coefficient between octanol and PBS buffer was determined. Radiolabelled compounds were dissolved in PBS to a concentration of approximately 9 μM and 50 μL of this solution was added to 450 μL PBS and 500 μL octanol into an eppendorf tube. The two phases were vigorously shaken for 20 min at 1400 rpm at room temperature (MS 3 basic vortexer, IKA, Staufen, Germany) followed by centrifugation for 2 min at 4500 rpm (Eppendorf Centrifuge 5424, Eppendorf AG, Hamburg, Germany).

Hereafter, 200 μL of each phase were collected and measured in a 2480 automatic Gamma counter Wizard 2 3” (PerkinElmer, Waltham, MA, USA). LogD value were calculated using Excel by dividing measured values of octanol by water and logarithmize the result. Values >0 reflects lipophilic-, values <0 hydrophilic compounds. (n = 3, six technical replicates)

###### Protein Binding

For this procedure siderophore conjugates were labelled as described before and diluted with PBS to a concentration of approximately 9 μM. 50 μL of this solution were added to 450 μL of PBS (control) or 450 μL of fresh human serum and incubated at 37°C for 30, 60 and 120 minutes. At each timepoint, 25 μL of PBS/serum were analyzed by size exclusion chromatography using MicroSpin G-50 columns (Sephadex G-50, GE Healthcare, Vienna, Austria) according to the manufacturer’s protocol. Hereafter, the column and eluate were measured separately in the gamma counter and percentage of protein bound conjugate was calculated by dividing measured counts of eluate by total counts. Radioactivity in the eluate reflects protein bound fraction and column bound, free labelled siderophore conjugate. (n = 3, three technical replicates)

###### Serum Stability

Serum stability probes were prepared according to protein binding section with PBS and fresh human serum. After 60, 120 and 240 minutes, 70 μL of serum/PBS were mixed with 70 μL of acetonitrile to precipitate proteins in the serum. Hereafter, the mixture was centrifuged for 1 minute and 70 μL of the supernatant were diluted with water and analyzed by radio-HPLC to determine intact labelled siderophore conjugate. (n = 2, two technical replicates)

###### Uptake and Competition Assay

Uptake assays were performed as previously described [17]. Briefly, 180 μL of *A. fumigatus* culture in iron-depleted and iron-replete (control – transporter suppression) aspergillus minimal media (AMM) [33] were added in pre-wetted 96-well MultiScreen Filter Plates HTS (1 μm glass fiber filter, Merck Millipore, Darmstadt, Germany) and incubated for 15 min at 37°C with either 25 μL PBS or 25 μL [Fe]TAFC solution (control – uptake block; ~10 μM). Hereafter, 50 μL of radiolabelled compound (final concentration approximately 90 nM) was added and incubated for another 45 min at 37°C. Hyphae were washed two times with ice-cold TRIS buffer and dry filters were measured in the gamma counter.

Competition assays were performed in the same way with slight modifications. Fungal cultures were pre-incubated with iron-labelled siderophore conjugates for 15 min (similar to blocking) and uptake values of [^68^Ga]Ga-TAFC into hyphae was determined after 45 min of incubation.

###### Growth Promotion Assay

This procedure was performed as previously described [34]. In this assay a mutant strain of *A. fumigatus* (*ΔsidA/ΔftrA*) that lacks the genes *sidA* and *ftrA* was used. These mutations impair both siderophore biosynthesis and reductive iron acquisition (RIA), in other words endogenous high-affinity iron acquisition [11], however this mutant is still able to take up siderophores. Consequently, utilization of siderophores can be simply tested by growth promotion assays allowing to analyze the effect of siderophore modification compared to the original molecule such as [Fe]TAFC In case of growth reduction distinction between reduced iron-utilization and antifungal activity is not possible. Conidia (10^4^) were point inoculated on solid 0.5 mL of iron depleted AMM in 24-well plates containing increasing concentrations of iron-labelled siderophore ranging from 0.1–50 μM. Plates were incubated for 48 h at 37°C in a humidity chamber and visually assessed[19]. (n = 3, three biological replicates)

###### Antifungal Susceptibility Assays

Antifungal susceptibility assays were performed with 96-well flat bottom plates (Greiner Bio-One GmbH) prepared with 100 μL of iron-depleted [Fe(-)] and -replete [Fe(+)] 2 x AMM containing 3*10^4^ spores of *A. fumigatus* (ATCC 46645) per well.

100 μL of antifungal siderophore conjugate, dissolved in water, was added to get a final concentration of 1 x AMM and siderophore conjugates in serial 2-fold dilutions in concentrations ranging from 256 - 0.016 μg/mL. Minimal inhibitory concentration (MIC) value was defined as the lowest concentration resulting in no visible fungal growth after 24h and 48h incubation at 37°C in a humidity chamber. Assays were repeated three times as biological replicates (n = 3).

Results were also displayed by taking microscopy pictures of each well after 24 h. Images were acquired with the IncuCyte S3 Live-Cell Analysis System equipped with a 20x magnification S3/SX1 G/R Optical Module (Essen BioScience Inc.). From each well a representative image was taken from the center of the well. Fungal growth was analyzed using the Basic Analyzer tool (Confluence %; Segmentation adjustment: 0; Adjust Size: 0) of the IncuCyte S3 software (Version 2019; Essen Bioscience Inc.). Images and confluence mask were exported in raw 8-bit images and raw 8-bit confluence mask, respectively. An overview of these pictures can be seen in the supplementary material. (Figure S3 and S4)

##### Animal experiments

All animal experiments were conducted in compliance with the Austrian Animal Protection laws and with approval of the Austrian Ministry of Science (BMWFW-66.011/0161-WF/V/3b/2016), the Czech Ministry of Education Youth and Sports (MSMT-21275/2016-2) and the institutional Animal Welfare Committee of the Faculty of Medicine and Dentistry of Palacky University in Olomouc.

###### In Vivo Stability and Ex Vivo Biodistribution

Stability test and biodistribution were conducted in 4-6 weeks old female BALB/c mice (in-house breed, ZVTA Innsbruck). ^68^Ga-labelled conjugates were injected via lateral tail vein using approximately 0.4 nmol of siderophore conjugate.

*In vivo* stability was determined by radio-HPLC analysis. After injection of the radiolabelled compound (~12 MBq) the mouse was euthanized after 10 minutes by cervical dislocation. Blood was collected and precipitated with ACN to remove proteins. Subsequently, the supernatant was diluted with water and used for further analysis. Urine samples were directly injected into the radio-HPLC. Percentage of intact radiolabelled siderophore conjugate was calculated by integration of the radiochromatogram (n=2) [35].

For biodistribution a similar procedure was applied but the mice were euthanized at 45 and 90 minutes. Hereafter, organs (blood, spleen, pancreas, stomach, liver, kidneys, heart, lung, muscle, femur) were removed and weighted. Samples were measured in a gamma counter and results expressed as percentage of injected dose per gram tissue (%ID/g). (n=3)

###### Invasive Pulmonary Aspergillosis Model in Rats

2-3 months old female Lewis rats were treated with the immunosuppressant cyclophosphamide (Endoxan, Bayter, 75 mg/kg i.p.) 5 days and 1 day before infected with *A. fumigatus*, to induce neutropenia. The animals received repeatedly (5 days, 1 day before and on the day of inoculation) antibiotic teicoplanin (Targocid, Sanofi, 35 mg/kg–5 days before i.m. or 25 mg/kg i.m.−1 day before and on the day of inoculation) to avoid bacterial superinfections and additional antibiotics were administered by drinking water (Ciprofloxacin, 2 mM, Polymyxin E, Colomycin, 0.1 mM) for the duration of the experiment. Infection in the lung was established by intratracheal inoculation of 100 μL of *A. fumigatus* spores (10^9^ CFU/mL *A. fumigatus* 1059 CCF) using TELE PACK VET X LED system equipped with a flexible endoscope (Karl Stroz GmbH & Co. KG, Tuttlingen, Germany) only [36].

###### PET/CT Imaging

*In vivo* PET/CT were conducted after 2-4 days, depending on the health condition of the animal and images were acquired with an Albira PET/SPECT/CT small animal imaging system (Bruker Biospin Corporation, Woodbridge, CT, USA). Radiolabelled antifungal siderophore conjugates were administered by retro-orbital (r.o.) injection of 5–10 MBq dose corresponding to ~2 μg of DAFC-conjugate per female Lewis rat.

Animals were anaesthetized with isoflurane (Forene^®^, Abbott Laboratories, Abbott Park, IL, USA) (2% flow rate) and positioned prone headfirst in the Albira system before the start of imaging. Static PET/CT imaging was carried out 45 min p.i. for all tested compounds. Infected animals were imaged three days after the inoculation with *A. fumigatus* spores. A 10-min PET scan (axial FOV 148 mm) was performed, followed by a triple CT scan (axial FOV 3 × 65 mm, 45 kVp, 400 μA, at 400 projections). Scans were reconstructed with the Albira software (Bruker Biospin Corporation, Woodbridge, CT, USA) using the maximum likelihood expectation maximization (MLEM) and filtered backprojection (FBP) algorithms. After reconstruction, acquired data were viewed and analyzed with PMOD software (PMOD Technologies Ltd., Zurich, Switzerland).

## 3. Results

### Synthesis and Radiolabelling

Precursor preparation of eflornithine by shielding the free amino groups with Boc-protection was obtained in moderate yield (>20%, non-optimized) but suitable for conjugation to [Fe]/[Ga]DAFC with standard HATU coupling. Synthesis of fludioxonil-butyric acid was carried out using a two-step reaction, which involved N-alkylation of the pyrrole with ethyl 4-bromobutyrate and followed by basic hydrolysis of ethyl ester group. Hereafter, the free carboxylic acid could easily couple to the [Fe]/[Ga]DAFC siderophore with very good yields over 60%. All other conjugates were directly linked by our standard HATU coupling strategy with yields of 20-80%. Exact analytical data are provided in the supplementary material.

Radiolabelling of all compounds was achieved with almost quantitative radiochemical yields (>95%) in 10 minutes at room temperature. Labelled compounds were used without any further purification.

#### *In Vitro* Characterization

##### LogD, Protein Binding and Serum Stability

Distribution coefficient (LogD), protein binding and serum stability in fresh human serum of all siderophore conjugates are summarized in Table 1. Log D values ranging from −3.45 to 1.30 revealed heterogeneous solubility properties of conjugated antifungals compared to LogD of the precursor molecule [Fe]DAFC with −2.34 [17].

**Table 1.**
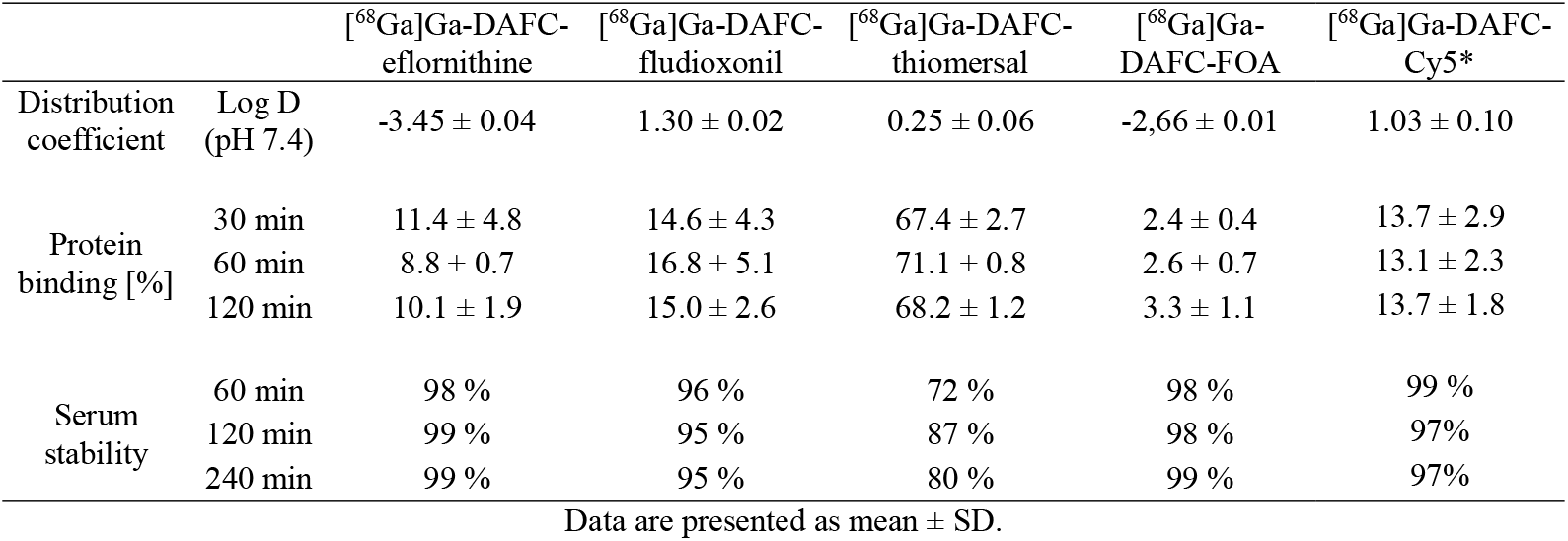
Distribution coefficient, protein binding and serum stability of siderophore compounds radiolabelled with gallium-68. Values of serum stability reflect percentage of intact radiolabelled conjugate. Results reflects three individual experiments; (*) Data from [19]

Protein binding revealed overall low and consistent values over time, except for [^68^Ga]Ga-DAFC-thiomersal which was ~70%. Compared to [^68^Ga]Ga-DAFC-FOA with an even higher LogD of 1.3 but lower protein binding (~4%), this could be explained by the reduced serum stability of [^68^Ga]Ga-DAFC-thiomersal which was ~80%.

##### Uptake and Competition Assay

Uptake and competition assays are summarized in Figure 2. Data of DAFC-conjugates are normalized to [^68^Ga]Ga-TAFC value of each experiments, respectively, to minimize biological variance. Uptake blocking by [Fe]TAFC results in a competition at the MirB transporter and should decrease the uptake. Under iron-replete conditions, MirB transporter is repressed at transcriptional level [37] and therefore the “uptake” under this condition represents unspecific uptake. All compounds showed reasonable values, i.e. although not as pronounced as found for [^68^Ga]Ga-TAFC, except for [^68^Ga]Ga-DAFC-Cy5 with very high unspecific binding in comparison to [^68^Ga]Ga-TAFC, which could not be reduced during blocking and iron-sufficient conditions. [^68^Ga]Ga-DAFC-fludioxonil revealed also higher values than [^68^Ga]Ga-TAFC but uptake reduction was possible by blocking and even more under iron-replete conditions indicating MirB-dependent uptake. [^68^Ga]Ga-DAFC-eflornithine, -thiomersal and -FOA showed a similar pattern, however, eflornithine unveiled only 20% uptake.

**Figure 2.**
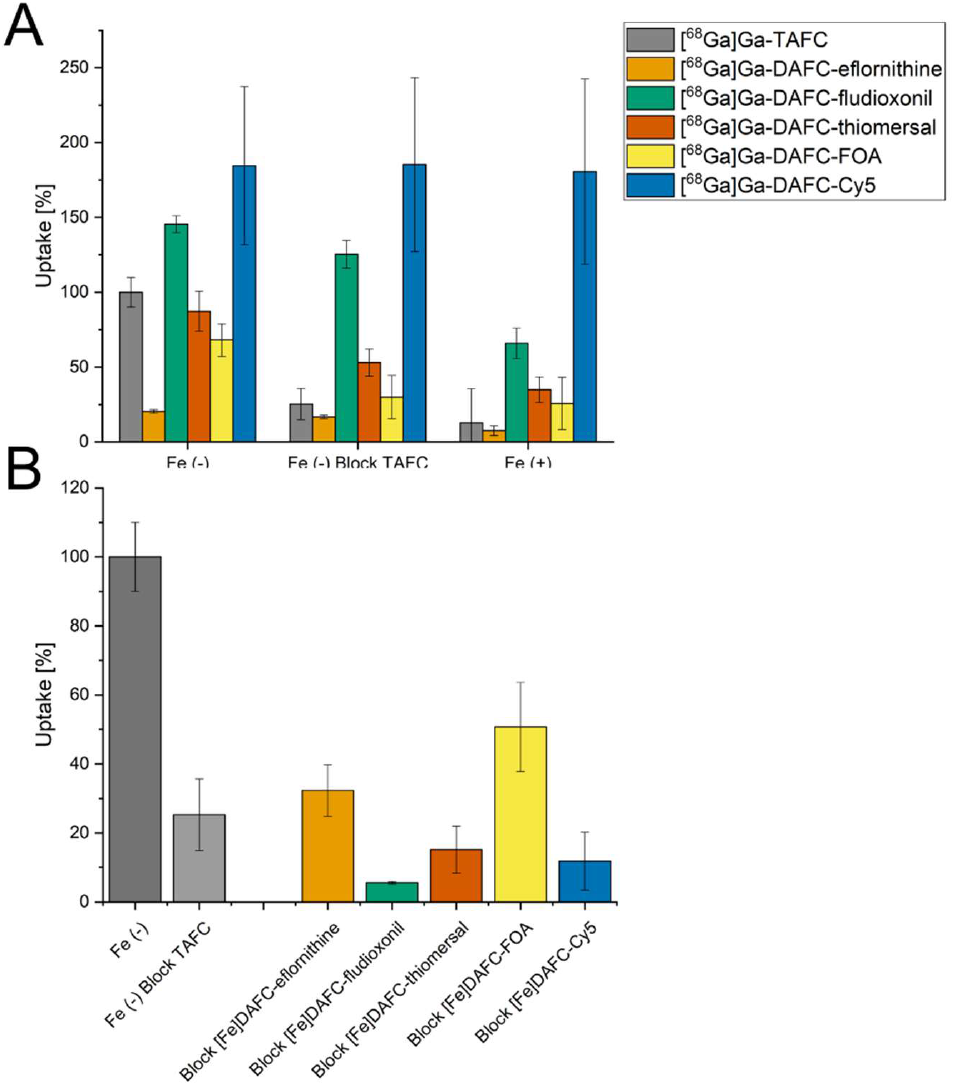
(A) Uptake of radiolabelled antifungal conjugates normalized to reference uptake of [^68^Ga]Ga-TAFC of each experiment, respectively. Grey bars represent control with [^68^Ga]Ga-TAFC. Blocking experiments with [Fe]TAFC due to competition at the MirB transporter should reduce uptake. Iron-replete conditions [Fe (+)] lead to a downregulation of MirB biosynthesis and active uptake of siderophores into the hyphae. (B) Competition assay of [^68^Ga]Ga-TAFC blocked with iron-containing antifungal conjugates in iron-depleted fungal culture. For all compounds a reduction of [^68^Ga]Ga-TAFC uptake could be observed, indicating a specific interaction with the MirB transporter. Data of DAFC-Cy5 adopted from reference [18].

In the ferric form, all conjugates were able to block uptake of [^68^Ga]Ga-TAFC in competition experiments, confirming interaction with the MirB transporter. All compounds showed comparable or even better blocking values compared to [Fe]TAFC.

##### Growth promotion assay

Growth promotion of the mutant strain of *A. fumigatus* (*ΔsidA/ΔftrA*) by [Fe]DAFC-conjugates is shown in Figure 3. Control with [Fe]TAFC showed a growth induction at 0.1 μM and sporulation at 10 μM, seen by the green colored conidia of *A. fumigatus*. Similar growth as in [Fe]TAFC containing media could be observed for [Fe]DAFC-eflornithine, -fludioxonil and -FOA, whereby with eflornithine and FOA conjugates sporulation was shown at 50 μM. Both, [Fe]DAFC-thiomersal and -Cy5 showed no growth promotion at all, indicating that iron from these conjugates cannot be utilized. It should also be considered that there is a competition between iron induced growth and a growth inhibitory effect of Cy5, but the positive impact of iron is only present if the fungus can utilize iron from the siderophore. This limitation should be kept in mind.

**Figure 3.**
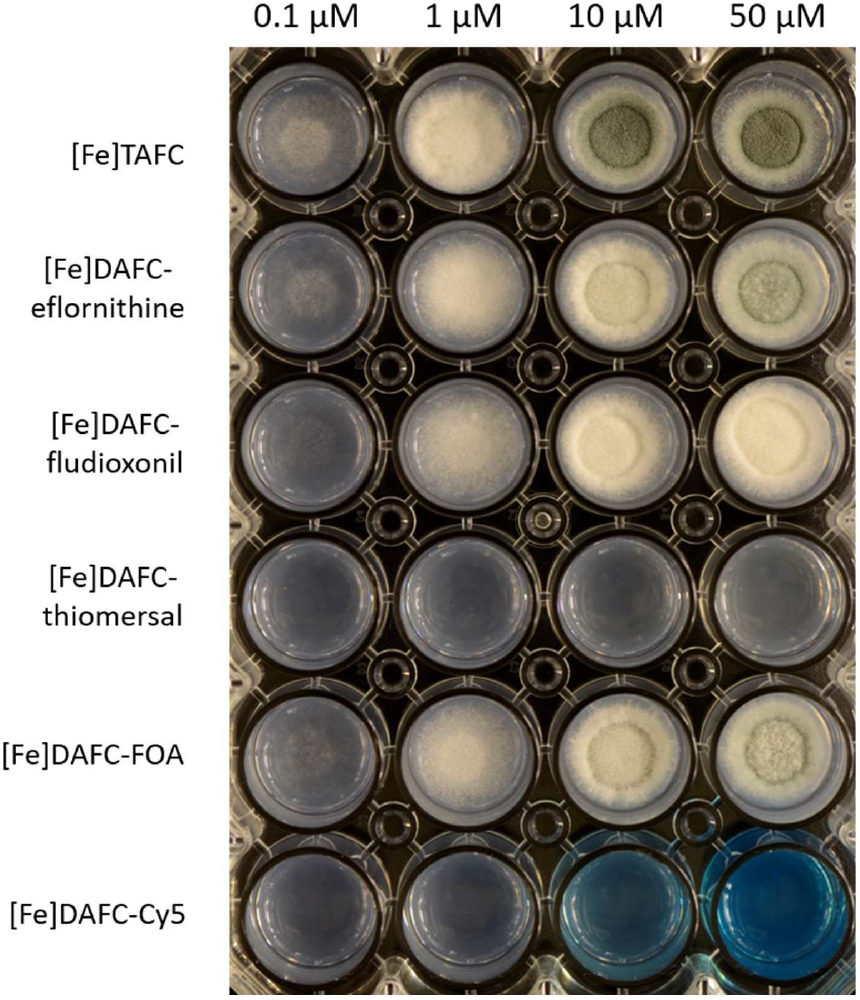
Growth promotion of *A. fumigatus* mutant strain Δ*sidA*/Δ*ftrA* after 48 h incubation at 37°C on iron depleted AMM with different concentrations of iron labelled conjugates. Hyphal growth can be distinguished from greenish (sporulation) and whitish (sterile) mycelia.

##### Antifungal susceptibility assays

Antifungal susceptibility assays were performed to evaluate the antifungal potential of the newly synthesized conjugates. Assays were performed as described in the method section, with defining the MIC as the lowest concentration resulting in no visible growth seen by the naked eye. A graphic overview is shown in Figure 4 and a list of all data is included in the supplementary material (Table S1). Most iron containing compounds showed a higher MIC value compared to their gallium labelled counterparts, except for DAFC-eflornithine and -thiomersal, which can be explained by the growth promoting effect of iron during iron starvation. Since gallium has an antifungal effect by itself [38], [Ga]TAFC is shown as a control but growth inhibition did not persist up to 48 h. Thiomersal conjugates showed the strongest growth inhibition but there was no difference to iron-sufficient media (MirB suppression) or to the original molecule, indicating that antifungal activity is not depending on active uptake by the fungus. [Ga]DAFC-fludioxonil, -FOA and -Cy5 showed promising MIC values, that were higher in iron-sufficient media (indicating MirB dependence) and lower than the original molecule (enhanced antifungal effect due to active uptake) except for fludioxonil. Chemical modification of the original molecule fludioxonil (0.5 μg/mL) reduced antifungal effect (fludioxonil butyric acid: 32 μg/mL see Table S1) but MIC value of the modified molecule was still preserved after conjugation. Eflornithine revealed no antifungal effect according to the definition of no visual growth, but still a growth reducing effect was observed. Microscopic pictures of all MIC results at 24 h can be seen in the supplementary material. (Figure S3 and S4)

**Figure 4.**
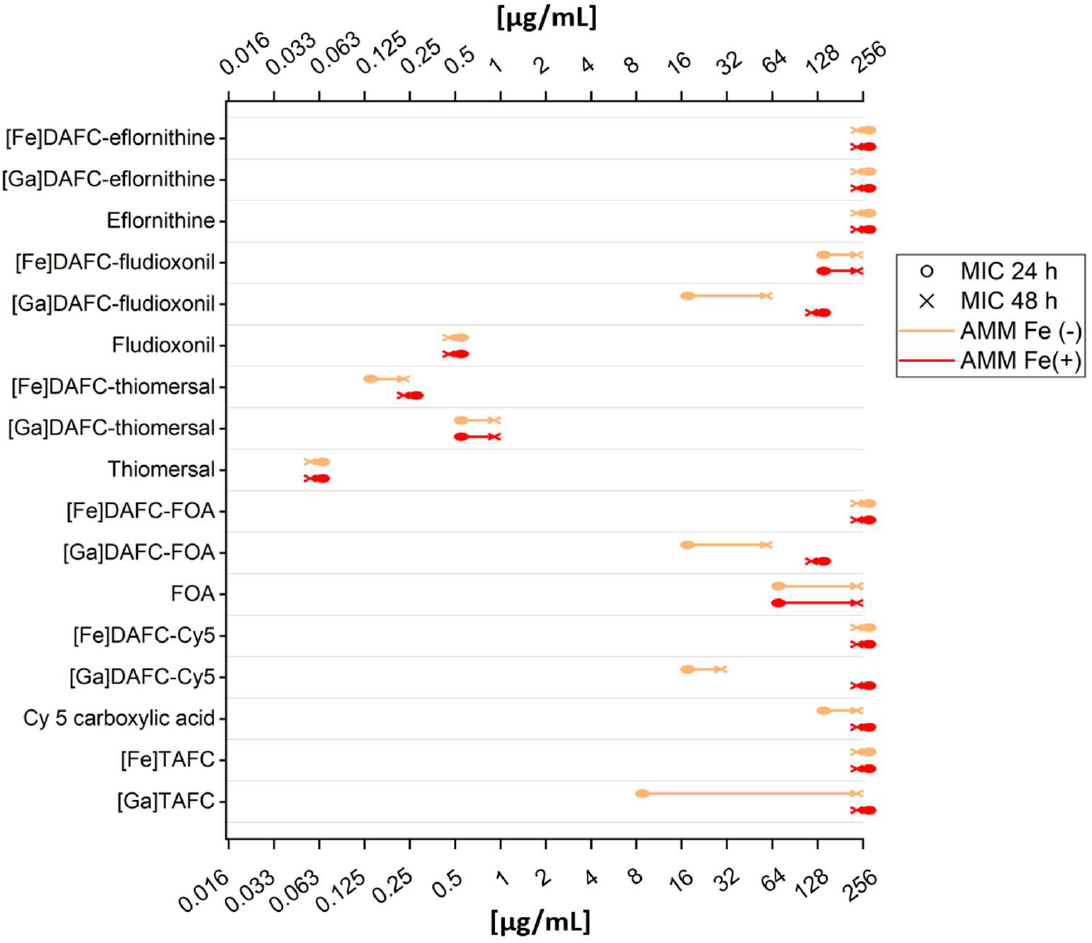
Scheme of all MIC results presented in this study. Orange lines present MIC values with iron-depleted medium (overexpression of the MirB transporter). Red lines are with iron-sufficient medium, which downregulates uptake by siderophore transporters such as MirB. Circle dot of the line displays 24 h and the “x” dot 48 h. Table with all data points is included in the supplementary material (Table S1).

#### *In vivo* Experiments

##### In vivo Stability and Biodistribution

Results of *in vivo* stability of the different compounds were very heterogeneous and shown in Table 2. [^68^Ga]Ga-DAFC-eflornithine and -FOA showed a very high stability both in blood and urine samples. [^68^Ga]Ga-DAFC-Cy5 showed no degradation in the blood but there was almost no intact conjugate after excretion through the urinary tract. The last two compounds [^68^Ga]Ga-DAFC-fludioxonil and - thiomersal revealed a high instability in both tested compartments. Only for thiomersal-conjugate instability was also be observed in the *in vitro* serum stability tests.

**Table 2.** *In vivo* stability of different antifungal conjugates radiolabelled with gallium-68. Samples form BALB/c mice 10 min after injection into the tail vain. Values reflect intact conjugate measured by radio-HPLC analysis. (Each value represents two biological replicates)

|  | [<sup>68</sup>Ga]Ga-DAFC-eflornithine | [<sup>68</sup>Ga]Ga-DAFC-fludioxonil | [<sup>68</sup>Ga]Ga-DAFC-thiomersal | [<sup>68</sup>Ga]Ga-DAFC-FOA | [<sup>68</sup>Ga]Ga-DAFC-Cy5 |
| --- | --- | --- | --- | --- | --- |
| Blood | 98.0% | 40.6% | 26.7% | >99.0% | >99.0% |
| Urine | 99.0% | 9.8% | 30.5% | >99.0% | 5.2% |

Biodistribution after 45 min and 90 min of antifungal conjugates are shown in Table 3. The main excretion of [^68^Ga]Ga-DAFC-eflornithine seemed to be the urinary tract with retention in the kidneys of around 20%, comparable to its high hydrophilicity and stability. On the contrary, [^68^Ga]Ga-DAFC-fludioxonil showed an accumulation on the intestine of the mouse and therefore indicated hepatobiliary excretion. Very high blood levels were found for [^68^Ga]Ga-DAFC-thiomersal, which correlated with the protein binding values found *in vitro*. Accumulation in the intestine and kidney retention was also found for this compound. Similar to these results [^68^Ga]Ga-DAFC-Cy5 showed overall higher blood levels but also an increasing accumulation of compound in the intestine (15-24%; 45-90 min) as well as in the kidneys (20-23%). This could be a limitation for PET imaging due to higher background signal. [^68^Ga]Ga-DAFC-FOA showed no significant accumulation in any organ over time.

**Table 3.** Biodistribution of gallium-68 labelled antifungal compounds in standard BALB/c mice. Injected into the tail vein and measured after 45 and 90 min, shown as injected dose per gram tissue (%ID/g). Important values are highlighted in bold. (Each value represents three biological replicates)

| Organ | <sup>68</sup>Ga]Ga-DAFC-eflornithine |  | <sup>68</sup>Ga]Ga-DAFC-fludiononil |  | <sup>68</sup>Ga]Ga-DAFC-thiomersal |  | <sup>68</sup>Ga]Ga-DAFC-FOA |  | <sup>68</sup>Ga]Ga-DAFC-Cy5 |  |
| --- | --- | --- | --- | --- | --- | --- | --- | --- | --- | --- |
|  | 45 min | 90 min | 45 min | 90 min | 45 min | 90 min | 45 min | 90 min | 45 min | 90 min |
| Blood | 0.93 ± 0.63 | 0.13 ± 0.03 | 0.17 ± 0.06 | 0.14 ± 0.02 | **10.35 ± 3.35** | **3.21 ± 0.94** | 1.30 ± 0.62 | 0.13 ± 0.04 | **4.38 ± 1.02** | **3.37 ± 0.65** |
| Spleen | 0.38 ± 0.05 | 0.22 ± 0.01 | 0.32 ± 0.06 | 0.30 ± 0.05 | 1.82 ± 0.64 | 0.61 ± 0.12 | 0.33 ± 0.08 | 0.12 ± 0.04 | 1.56 ± 0.15 | 5.58 ± 0.94 |
| Pancreas | 0.31 ± 0.07 | 0.06 ± 0.01 | 0.05 ± 0.01 | 0.04 ± 0.00 | 1.59 ± 0.61 | 0.45 ± 0.09 | 0.28 ± 0.04 | 0.06 ± 0.02 | 0.56 ± 0.07 | 0.46 ± 0.12 |
| Stomach | 0.28 ± 0.06 | 0.09 ± 0.03 | 0.94 ± 1.12 | 0.25 ± 0.22 | 1.77 ± 0.50 | 0.72 ± 0.24 | 0.40 ± 0.07 | 0.13 ± 0.07 | 3.95 ± 3.13 | 1.77 ± 1.61 |
| Intestine | 0.68 ± 0.08 | 0.51 ± 0.05 | **23.08 ± 2.43** | **25.19 ± 1.18** | **19.41 ± 7.26** | **13.63 ± 1.37** | 2.68 ± 0.54 | 2.52 ± 0.40 | **15.82 ± 0.31** | **24.73 ± 0.34** |
| Kidneys | **23.01 ± 2.23** | **19.35 ± 4.55** | 0.22 ± 0.03 | 0.20 ± 0.03 | **7.43 ± 2.24** | **3.67 ± 0.98** | 2.72 ± 0.21 | 1.75 ± 0.21 | **4.60 ± 0.52** | **3.37 ± 0.68** |
| Liver | 0.55 ± 0.03 | 0.38 ± 0.05 | 2.20 ± 0.49 | 0.83 ± 0.23 | 3.67 ± 0.38 | 2.37 ± 0.83 | 0.70 ± 0.34 | 0.60 ± 0.38 | **19.73 ± 2.25** | **22.95 ± 1.57** |
| Heart | 0.28 ± 0.07 | 0.07 ± 0.01 | 0.07 ± 0.02 | 0.06 ± 0.02 | 3.03 ± 0.91 | 0.97 ± 0.14 | 0.33 ± 0.09 | 0.07 ± 0.00 | 1.39 ± 0.11 | 1.25 ± 0.25 |
| Lung | 0.89 ± 0.16 | 0.24 ± 0.04 | 0.32 ± 0.26 | 0.18 ± 0.09 | 7.04 ± 2.67 | 2.96 ± 1.75 | 0.66 ± 0.02 | 0.21 ± 0.06 | 2.95 ± 0.20 | 1.95 ± 0.25 |
| Muscle | 0.33 ± 0.23 | 0.04 ± 0.01 | 0.04 ± 0.01 | 0.03 ± 0.01 | 1.25 ± 0.49 | 0.36 ± 0.10 | 0.20 ± 0.02 | 0.05 ± 0.02 | 0.50 ± 0.12 | 0.45 ± 0.11 |
| Femur | 0.21 ± 0.09 | 0.11 ± 0.04 | 0.07 ± 0.02 | 0.09 ± 0.06 | 1.50 ± 0.72 | 0.56 ± 0.22 | 0.45 ± 0.17 | 0.14 ± 0.03 | 0.80 ± 0.43 | 0.85 ± 0.13 |

Data are presented as mean ± SD.

##### PET/CT images

In Figure 5 coronal PET/CT slices of non-infected Lewis-rats are shown, injected with different ^68^Ga-labelled antifungal siderophore conjugates. [^68^Ga]Ga-DAFC-eflornithine and -FOA showed a primary excretion through the urinary tract with clear delineation of the kidneys and bladder. As already described in the biodistribution experiments, [^68^Ga]Ga-DAFC-fludioxonil, -thiomersal and -Cy5 were excreted through the hepatobiliary way which resulted in a high signal in the intestinal region.

**Figure 5.**
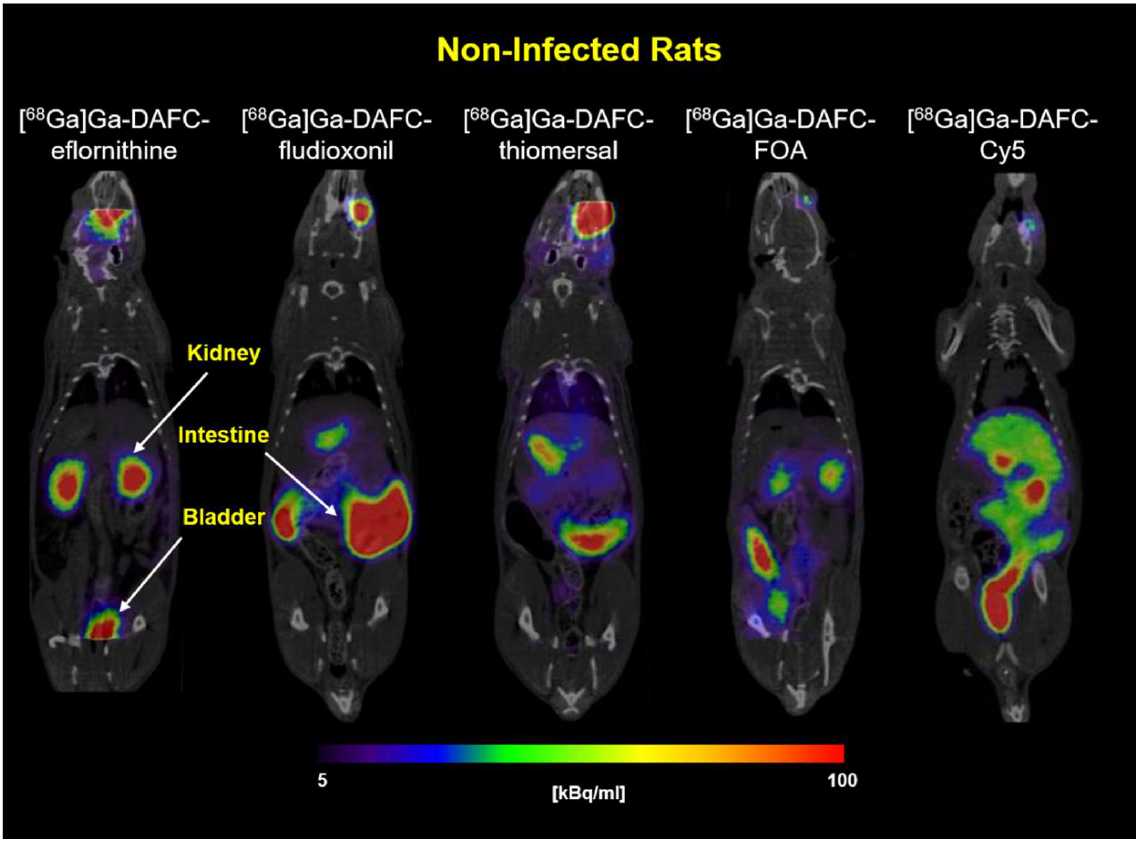
Coronal PET/CT of non-infected Lewis-rats after 45 min post injection of 68Ga-labelled siderophore conjugates (approx. 5–10 MBq injected dose). Kidneys, intestine and bladder are highlighted by arrows in the picture. Radioactive spots in the eye region originate from the retro-orbital injection.

PET/CT images of Lewis-rats infected with *A. fumigatus* in the lung, displayed an accumulation of all antifungal conjugates in the infected region. (Figure 6) CT images of the lung revealed anatomical changes with co-localization of radioactive signal. Skriba et al. showed that these anatomical changes originate from infection with *A. fumigatus* which was also confirmed by histological examination [36]. Compared to non-infected animals, no significant radioactive signal could be observed in the lung region for all siderophore conjugates.

**Figure 6.**
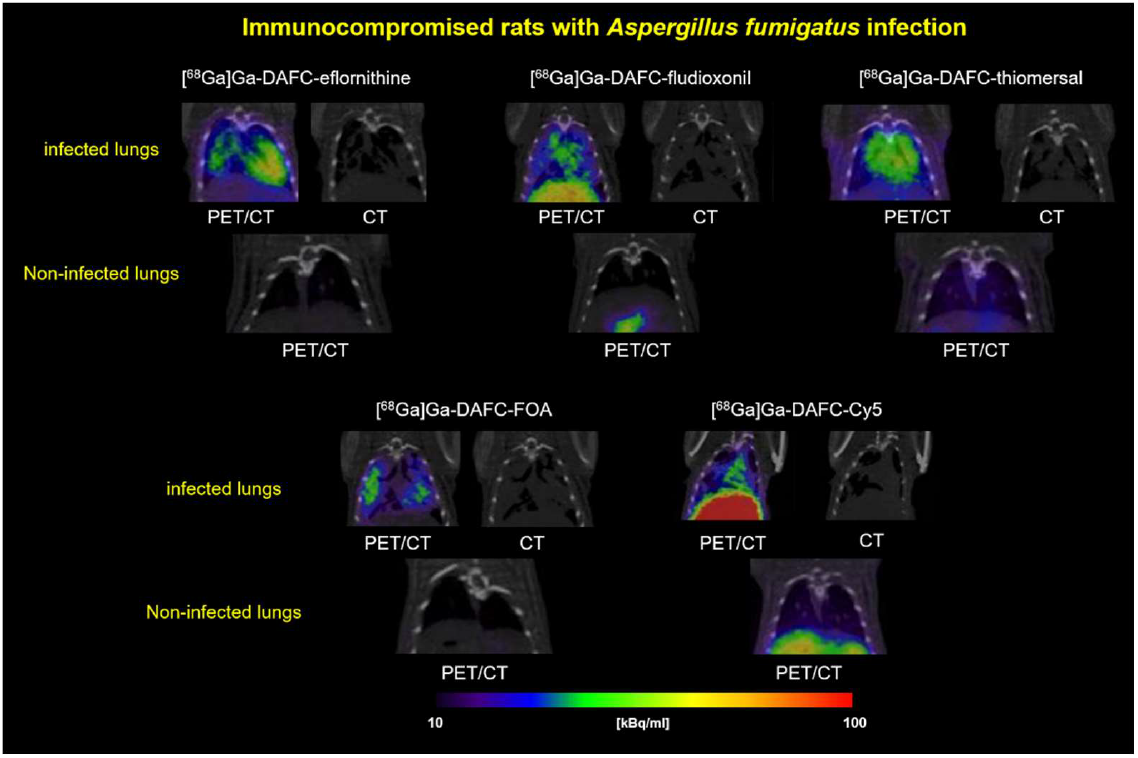
Coronal PET/CT slices of immunocompromised Lewis-rats infected with *A. fumigatus* in the lung. Pictures are showing the lung section of infected (top row) and non-infected (bottom row, control) rats of each compound respectively. Animals were injected retro-orbitally and pictures made after 45 min with approx. 5-12 MBq injected dose. CT images were added to show the severity of the infected lung tissue.

## 4. Discussion

Antimicrobial siderophores have been attracting the interest of scientists for many decades. First discovered as natural products of bacteria, so called sideromycins, they were investigated as antibacterial compounds e.g. salmycin and albomycin for growth inhibition of *Streptococcus pneumoniae* and *Staphylococcus aureus* [39,40]. This natural concept of utilizing the iron acquisition system of bacteria to channel antibacterial molecules inspired the development of new synthetic compounds. For example, pyoverdine (siderophore) conjugated with ampicillin (beta-lactam antibiotic) against *Pseudomonas aeruginosa* [41]. The big advantage of the synthetic strategy is the high variability of conjugation partners. It is possible to use these siderophores as a “Trojan horse” to smuggle all kinds of molecules into bacteria [42]. The recently approved drug Cefiderocol (Fetroja^®^) by the FDA (Nov 2019) is the first artificial sideromycin on the market consisting of a catechol chelating unit and a cephalosporin antibiotic (beta-lactam) against gram-negative bacteria [43].

This Trojan horse concept can be adopted for fungal species as well, using fungal specific siderophores to deliver antifungal molecules. Previous research published by our group showed that fluorescence dyes could be attached to [Fe]DAFC and a specific fluorescence signal could be detected in *A. fumigatus* hyphae that was dependent on active uptake via MirB [18]. Furthermore, “large” antifungal peptides were attached to [Fe]DAFC to inhibit fungal growth but unfortunately no antifungal effect could be observed [34]. In this study we attempted to modify TAFC with small organic molecules that have known antifungal properties described in the literature and label them with gallium-68 to perform PET/CT images of *A. fumigatus* lung infections in a rat model.

Synthesis of the conjugates was straight forward, with high purity of the resulting compounds. Modifications had to be made for eflornithine, to provide better synthetic yields with a lower rate of side products. Fludioxonil had to be chemically modified by adding a linker function to its molecular structure for coupling to DAFC. The new compounds were labelled with gallium-68 to perform *in vitro* and *in vivo* tests. Labelling resulted in a high radiochemical purity under mild conditions (10 min, room temperature, pH = 4.5).

Siderophore conjugates used within this research were produced in two different modifications: iron-containing ([Fe]DAFC) and gallium-labelled ([Ga]DAFC) conjugates. The rationale behind using gallium instead of iron was to enhance targeting efficiency because the iron transported by siderophores leads in *A. fumigatus* after uptake to downregulation of siderophore transporters such as MirB [44]. This would lead to a reduction or even cessation of siderophore uptake into the hyphae and in the case of antifungal conjugates, resulting in a better survival of the fungus. Additionally, iron as an essential nutrient has a growth-enhancing effect on fungi. Gallium nitrate is a FDA approved drug, used in cancer-related hypercalcemia (Ganite^®^) and showed a very good tolerance in patients [45]. It should be considered, that gallium itself has an antifungal effect on microorganisms [46]. Bastos et al. have shown that Ga(NO3)3 has an antifungal effect on *A. fumigatus* cultures in different media [38] and Ga(NO3)3 is also currently investigated against *Pseudomonas aeruginosa* lung infection of adults with cystic fibrosis in a clinical trial [47]. So far, it is not completely clear how gallium influences the antifungal activity of the described siderophore conjugates but MIC tests with [Ga]TAFC showed an inhibition of growth at 8 μg/mL and 24 h but after 48 h no inhibition was observed anymore. Fungistatic behavior of Ga(NO3)3 have also been described by Bastos et al. in AMM [38]. As seen in Figure 4, an overall better fungal growth inhibition was observed for gallium labelled conjugates in comparison to iron labelled counterparts. In the development of antimicrobial siderophore conjugates as “Trojan horse”, Miller, Nolan, and others, who generated fundamental knowledge in antibiotic conjugates, always used iron for labelling of their compounds [48–51]. One of the first reports on enhancing antibacterial potency by labelling with gallium was published by Pandey et al. with Ferrichrome-Ciprofloxacin against *Escherichia coli*. They observed a 2-4 fold enhancement of activity compared to the iron complex [52].

Antifungal susceptibility assays revealed the potential of fungal growth inhibition by the produced antifungal siderophore conjugates. To test their MirB transporter dependency, two different media were used: iron depleted AMM Fe(-) (upregulation of MirB transporter,) and iron sufficient media AMM Fe(+) (control, no active transport). Except for [Ga]DAFC-eflornithine and –thiomersal, all gallium-labelled compounds showed a MirB dependent antifungal effect. Thiomersal conjugates, revealed an overall low MIC value independent on the media and labelling with gallium or iron, which is probably due to cleavage outside the hyphae resulting in ethylmercury (active metabolite) causing growth inhibition. Higher instability in human serum and *in vivo* fortifies these assumption but this limits the applicability, because adverse health effects of ethylmercury are still controversially discussed [31]. [Ga]DAFC-fludioxonil (16 μg/mL), -FOA (16 μg/mL) and Cy5 (16 μg/mL) showed a growth reduction of 3 to 4 dilution steps in iron depleted media compared to iron sufficient after 24 h giving a proof of principle for a MirB dependent antifungal effect. eflornithine revealed no antifungal effect independent of the siderophore conjugation.

Competition assay with iron-containing antifungal conjugates blocking uptake of [^68^Ga]Ga-TAFC showed a specific interaction of all conjugates with the MirB transporter encouraging the hypothesis of a transporter dependent effect. Uptake assays of [^68^Ga]Ga-DAFC-fludioxonil, -thiomersal, -FOA and - Cy5 showed a high accumulation by *A. fumigatus* hyphae, however, uptake of Cy5 could not be blocked by [Fe]TAFC or iron sufficient media. This unspecific binding was already observed in earlier studies and origins probably due to interaction with the outer cell compartment [18,19]. Uptake of [^68^Ga]Ga-DAFC-eflornithine was notably low, which could be one reason, why no antifungal effect was observed when coupled to [Ga]DAFC. Growth assays confirmed utilization of DAFC-eflornithine, -fludioxonil, and FOA, whereas [Fe]DAFC-Cy5 and -thiomersal did not promote growth, providing limited additional information towards the theranostic properties of these compounds.

Beside growth inhibiting properties, pharmacokinetics of these novel antifungal siderophore conjugates should be especially considered, which could predict diagnostic and clinical properties. [53] PET/CT images of non-infected rats show the distribution of radioactive labelled compound and main excretion routes supported by biodistribution results. [^68^Ga]Ga-DAFC-eflornithine and FOA were manly excreted through the kidney and bladder of the rats. These findings are in line with a low protein binding capacity and a hydrophilic character of the compounds. In a diagnostic setting, a rapid elimination from the body lowers radiation burden for the patient and allows high image resolution. Lipophilic compounds like [^68^Ga]Ga-DAFC-fludioxonil, -thiomersal and -Cy5 are manly excreted through liver and intestine as seen in PET/CT images and in biodistribution studies. This would lead to a higher background signal, which would be less of a problem since IPA infections are located in the lungs, but it would lead to a higher radiation burden to the patient. Increased blood levels of [^68^Ga]Ga-DAFC-thiomersal and -Cy5 would be also not beneficial from a diagnostic prospective, however, a therapeutic effect could be enhanced due to longer circulation times. Studies from the recently approved drug Cefiderocol (Fetroja^®^) against gram-negative bacteria revealed an *in vitro* plasma binding of 58 % and a main clearance through the kidneys (>90%) [54]. A longer circulation time of antifungal siderophore compound could increase the likelihood of uptake by the fungus resulting in a better therapeutic outcome.

Diagnostic applicability of these compounds was tested in an *A. fumigatus* lung infection in a rat model. All conjugates revealed uptake into the infected area of the lung and a visualization in PET/CT. This specific accumulation holds promise for a therapeutic effect *in vivo*. Images were similar to previous infections images with [^68^Ga]Ga-TAFC by Petrik et al. [55] and modified [^68^Ga]Ga-TAFC by Kaeopookum et al. [17]. PET/MRI pictures by Henneberg et al. using the *Aspergillus* specific antibody hJF5 labelled with copper-64 to visualize an *A. fumigatus* infection, showed also similar patterns in a mouse model [56].

Taken together, antifungal siderophore conjugates synthesized within this study are highly interesting model compounds for theranostic applications in fungal infections. The concept of using siderophore conjugates for theranostics, combining gallium-68 for PET with a therapeutic moiety have first been described by Ferreira et al [57]. They coupled siderophore catecholate binding moieties to a DOTAM scaffold, and additionally ampicillin for antibacterial activity. The iron containing compound did not reveal antibacterial activity in wild type *E.coli*, but specific *in vitro* uptake could be proven. *In vivo* data of the therapeutic conjugate have not been provided. More recently Pandey et al. [52] reported a siderophore ciprofloxacin conjugate based on ferrichrome. They showed a siderophore iron transport-dependent internalization of one compound and antibacterial activity against *Peudomonas aeruginosa, Staphylococcus aureus* and *Klebsiella pneumoniae* of the Gallium (III) complex, whereas the Ferric complexes remained inactive, similar to our findings. They also reported biodistribution data in normal mice, but no data in animal infection models. Both studies also failed to provide proof of a therapeutic effect in respective infection models. Our study overall confirmed these findings for theranostics in fungal disease, which so far have not been described. Additionally we provided for the first time, specific *in vivo* targeting of infections with such “Trojan horse” conjugates by PET imaging and could show that the antifungal efficacy in most compounds was clearly related to the siderophore transport target MirB. Data obtained in this study give highly valuable information for synthesis of future compounds with higher antifungal potency which bear great potential for diagnostic and therapeutic use.

## 5. Conclusions

In this study, we successfully modified TAFC with a variety of antifungal molecules with divergent mechanism of action, unrelated to commonly used therapeutics. Targeting the iron acquisition system of *A. fumigatus* with these antifungal siderophore conjugates revealed a proof of principle to utilize MirB for delivering a variety of antifungals, to inhibit fungal growth in antifungal susceptibility tests. By labelling the compounds with gallium-68, a lung infection of *A. fumigatus* in a rat model was confirmed, also revealing insights about the influence of the conjugation partners on pharmacokinetic properties. Knowledge gained within this research could lead to sophisticated strategies in the field of IPA infections, combining imaging and therapy in a theranostic approach.

## Supporting information

Supplement_Antifungal_Pfister_2021

## Abbreviation

*A. fumigatus*: *Aspergillus fumigatus*
AMM: Aspergillus minimal medium
ACN: Acetonitrile
DAFC: Diacetylfusarinine C
FsC: Fusarinine C
HATU: O-(7-Azabenzotriazol-1-yl)-N,N,N’,N’-tetramethyluronium hexafluorophosphat
MIC: Minimal inhibitory concentration
Na_2_EDTA: Ethylenediaminetetraacetic acid disodium salt
PBS: Phosphate buffer saline
SIT: Siderophore iron tranpsorter
TAFC: Triacetylfusarinine C
TRIS: Tris(hydroxymethyl)aminomethane
PET/CT: Positron Emission Tomography/Computer Tomography

