## Supplement_Antifungal_Pfister_2021 for "Antifungal siderophore conjugates for theranostic applications in invasive pulmonary aspergillosis using low molecular TAFC scaffolds"

**MALDI-TOF MS:** Matrix-assisted laser desorption/ionization time -of-flight mass spectrometry was performed on a Bruker microflex™ bench-top MALDI-TOF MS (Bruker Daltonics, Bremen, Germany). Samples were prepared on a microscout target (MSP96 target ground steel BC, Bruker Daltonics) using dried-droplet method and α-cyano-4-hydroxycinnamic acid (HCCA, Sigma-Aldrich, Handels GmbH, Vienna, Austria) as matrix. All spectra were recorded by summarizing 600 laser shots per spot and Flex Analysis 2.4 software was used for data processing.

**$^1$H-NMR Spectroscopy**

$^1$H-NMR spectra were recorded on "Mars" 400 MHz Bruker Avance 4 Neo spectrometer in $CDCl_3$ or DMSO-d6, respectively. Chemical shifts are expressed in ppm downfield relative to tetramethylsilane and the coupling constants (J) are reported in Hertz. Data for $^1$H-NMR spectra are reported as follows: s = singlet, br s = broad singlet, d = doublet, dd = doublet of a doublet, t = triplet, q, quartet, m = multiplet.

**Analytical RP-HPLC:** Reversed-phase (RP) high-performance Liquid chromatography (HPLC) analysis was carried out using the following instrumentation: UltiMate 3000 RS UHPLC pump, UltiMate 3000 autosampler, UltiMate 3000 Variable Wavelength Detector; UV detection at λ = 220nm (Dionex, Germering, Germany) Radio-detector (Gabi Star, Raytest; Straubenhardt, Germany) using Jupiter 5 µM $C_{18}$ 300 Å 150 x 4.6 mm (Phenomenex Ltd. Aschaffenburg, Germany) as column with acetonitrile (ACN)/$H_2O$/0.1% trifluoroacetic acid (TFA) as mobile phase; flow rate of 1 mL/min;

**Gradient A:** 0.0–3.0 min 10% ACN, 3.0–16.0 min 10–60 % ACN, 16.0–18.0 min 60% ACN, 18.0–18.1 min 60–10% ACN, 18.1–22.0 min 10% ACN.

**Gradient B:** 0.0–3.0 min 10 % ACN, 3.0–16.0 min 10–100 % ACN, 16.0–18.0 min. 100 % ACN, 18.0-18.1 min 100-10 % ACN, 18.1–23.0 min. 10 % ACN

**Preparative RP-HPLC.** Sample purification via RP-HPLC was carried out on a Gilson 322 Pump with a Gilson UV/VIS-155 detector (UV detection at λ = 220 nm) using a PrepFC™ automatic fraction collector (Gilson, Middleton, WI, USA), Eurosil Bioselect Vertex Plus 30 x 8 mm 5 µm $C_{18}$A 300Å pre-column and Eurosil Bioselect Vertex Plus 300 x 8 mm 5 µm $C_{18}$A 300Å column (Knauer, Berlin, Germany) and following ACN/$H_2O$/ 0.1 % TFA gradients with a flow rate of 2 mL/min:

**Gradient 1**: 0.0–1.0 min 0 % ACN, 1.0–35.0 min 0–50 % ACN, 35.0–36.0 min 50 % ACN, 36.0–36.1 min 50–0 % ACN, 36.1–43.0 min 0 % ACN

**Gradient 2**: 0.0–1.0 min 20 % ACN, 1.0–36.0 min 20–100 % ACN, 36.0–40.0 min 100 % ACN, 40.0–40.1 min. 100–20 % ACN, 40.1–47.0 min. 20 % ACN,

**Gradient 3:** 0.0–1.0 min 10% ACN, 1.0–36.0 min 10–80 % ACN, 36.0–37.0 min 80 % ACN, 37.0–37.1 min 80-10% ACN; 37.1–43.0 min 10% ACN

**Gradient 4:** 0.0–1.0 min 10 % ACN, 1.0–35.0 min 10–50 % ACN, 35.0–36.0 min 50 % ACN, 36.0–36.1 min 50–10 % ACN, 36.1–43.0 min 10 % ACN

#### Precursor preparation:

[Fe]Fusarinin C ([Fe]FsC)/[Ga]FsC): Fusarinin C was obtained by fungal culture according to Schrettl *et al* [1]. *Aspergillus fumigatus* mutant strain *ΔsidG* (which lacks the enzyme for acetylation of Fusarinine C) was grown ($1*10^6$ Spores/mL) in 200 mL iron depleted minimal medium, incubated for 28 h at 37°C and shaking at 200 rpm. After filtering of the culture supernatant, $FeSO_4$ was added to a final concentration of 10mM to get a red coloured solution. To produce [Ga]FsC, $GaBr_3$ was used instead of the $FeSO_4$, resulting in a colourless solution. The filtrate was subsequently loaded to a FlashPure cartridge (C18; 40 µm; 12g; column volume (CV) of 24 mL; BÜCHI Labortechnik AG, Flawil, Switzerland) by using a REGLO tubing pump (Type ISM795, Ismatec SA, Glattbrugg-Zürich, Switzerland) with a flow rate of 10 mL/min. Fixed [Fe]/[Ga]FsC on the cartridge was washed with 2 CV of water and then eluted with 5 CV of methanol. After evaporation of the organic solvent approximately 250 mg of [Fe]FsC could be obtained as a red-brown coloured solid with a purity of >90% confirmed by analytical RP-HPLC (gradient A $t_R$ = 9.0 min) and the product was used for synthesis without further purification. MALDI-TOF-MS: *m/z* [M+H] = 780.68 [$C_{33}H_{51}FeN_6O_{12}$; exact mass: 779.63 (calculated)]. [Ga]FsC resulted in a yellow coloured solid (gradient A $t_R$ = 8.7 min) MALDI-TOF-MS: *m/z* [M+H] = 793.68 [$C_{33}H_{51}GaN_6O_{12}$; exact mass: 792.28 (calculated)]

**Acetylation of [Fe]/[Ga]FsC:**

To acetylate FsC, 20-30 mg (38 µmol) dissolved in 500 µL water was rocked with 10-20 µL (0.2 µmol) of acetic anhydride for 2 min at room temperature and intense shaking. Resulting products mono-, di-, and triacetylfusarinine C were immediately purified via preparative RP-HPLC using gradient 1 to collect N,N´-diacetylfusarinine C ([Fe]/[Ga]DAFC, $t_R$ = 17.8/18.1 min) in high purity (> 95%) followed by lyophilization. MALDI-TOF-MS: $t_R$ ([Fe]DAFC= 10.9 min *m/z* [M+H] = 864.01 [$C_{37}H_{55}FeN_6O_{14}$; exact mass: 863.70 (calculated)] [2]; [Ga]DAFC was produced in the same way except of using [Ga]FsC as a starting material. Analytical data: $t_R$ ([Ga]DAFC= 10.7 min *m/z* [M+H] = 877.37 [$C_{37}H_{55}GaN_6O_{14}$; exact mass: 876.30 (calculated)]

**Modification of Fludioxonil**

NaH (60% dispersion in mineral oil, 0.69 g, 17.2 mmol) were prewashed with pentane to get a white powdered dispersed in 30 mL DMF. Fludioxonil (2.01 g, 8.0 mmol) was dissolved in DMF (40 mL) and added dropwise to the NaH dispersion under argon atmosphere and stirred for 2h at room temperature. NaI (0.35 g, 2.3 mmol) and ethyl 4-bromobutyrate (2.5 g, 12.8 mmol) in DMF (30 mL) were added and stirred for additional 3 h. Hereafter the mixture was poured in ice cold water (250 mL) and extracted with ether. Combined organic layers were washed with brine and dried over $Na_2SO_4$. The solvent was removed under vacuum and the residue chromatographed with normal phase column, using $CH_2Cl_2$/ethyl acetate (1:2) to give the ester compound (3.04 g, 8.4 mmol: >100 % yield with some impurities) as white crystals. [3] $^1$NMR (400 MHz, $CDCl_3$) δ (ppm): 7.72 (1H, d, J= 7.6 Hz), 7.26 (1H, br s), 7.16-7.12 (2H, m), 6.97 (1H, d, J= 7.6 Hz), 4.16 (2H, q, J= 6.7 Hz, $OCH_2$), 4.05 (2H, t, J= 6.6 Hz, $NCH_2$), 2.33 (2H, t, J= 6.2 Hz, $CH_2COOEt$), 2.19-2.12 (2H, m, $CH_2$), 1.27 (3H, t, J= 6.7 Hz, $CH_3$).

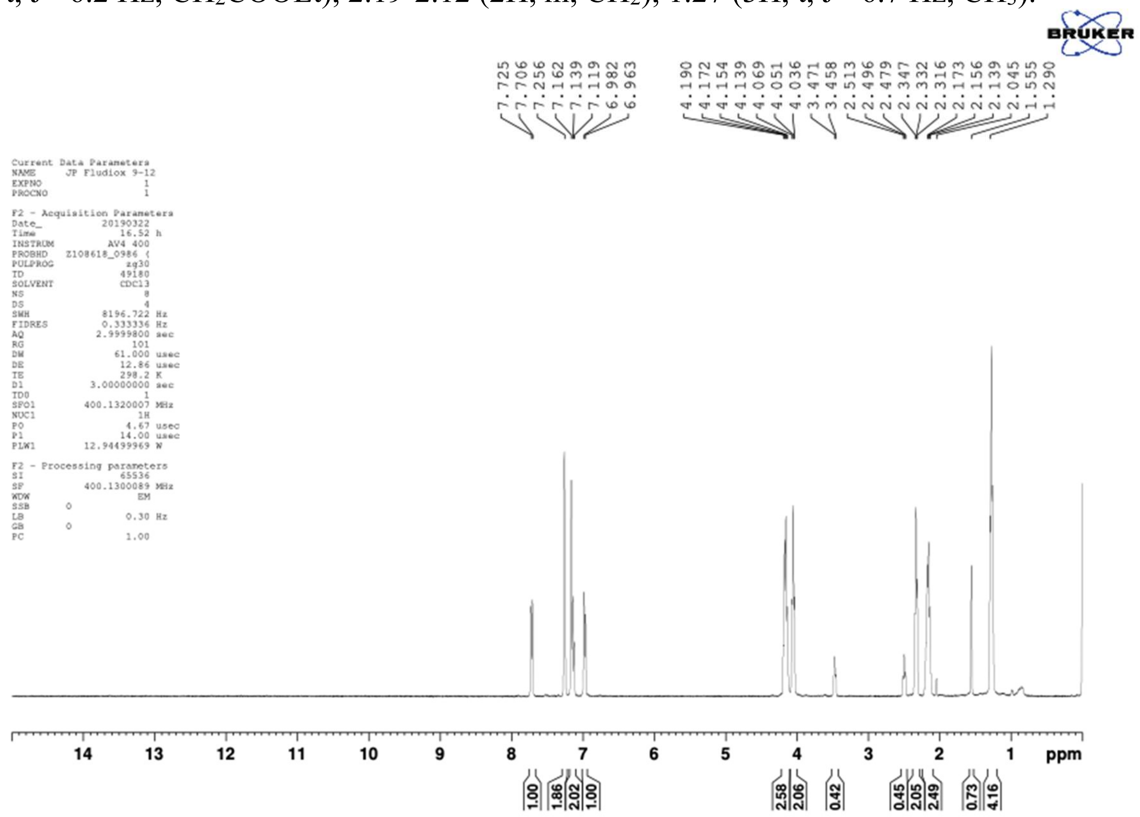


**Figure S1.** $^1$H-NMR spectrum of ethyl fludioxonil-butyrate.

For cleavage of the ester group, 13.5 mL (16.7 µmol) of a 5% NaOH solution in water was added to the product dissolved in ethanol (40 mL) and the mixture was stirred for 2h at room temperature. Reaction solution was acidified in the ice bath with 2N HCl to adjust the pH to 1-2. Hereafter, the organic solvent was removed under vacuum and the precipitate filtered and dried to give the final product (1.156 g, 3.46 mmol, yield of 51 %) as a white powder. [4] NMR (400 MHz, DMSO) δ (ppm): 7.85 (1H, d, J=2.4 Hz), 7.52 (1H, dd, J=8.0 Hz, 1.6 Hz), 7.41 (1H, d, J = 2.4 Hz), 7.34-7.27 (2H, m), 4.04 (2H, t, J = 7.0 Hz, N-$CH_2$), 2.20 (2H, t, J = 7.4 Hz, $CH_2$-COOH), 2.01-1.94 (2H, m, $CH_2$).

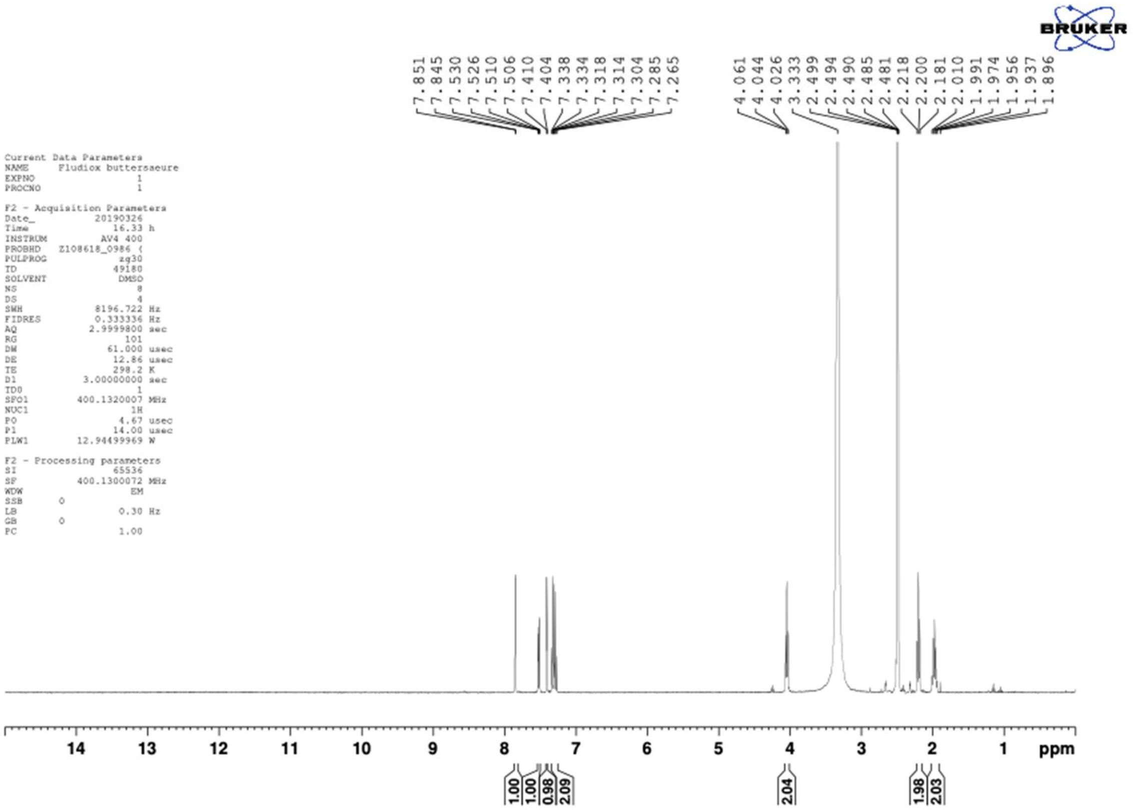


**Figure S2.** $^{1}$H-NMR spectrum of fludioxonil-butyric acid

### Conjugation of the antifungals:

##### Conjugation of Eflornithine

Eflornithine contains two primary amines that could interfere during the conjugation process and produce polymers. Therefore, as a first step, the free amine groups had to be protected with tert-butyloxycarbonyl protecting group (Boc). 100 mg (423 µmol) of eflornithine were dissolve in 750 µL water, mixed with 230 mg (1.05 mmol) of Boc-anhydride, dissolved in 500 µL dioxan, and left to react for 24h at room temperature. End of the reaction was confirmed by a Kaiser-Test and product was dried under vacuum and dissolved again in water. By adding 40 µL of NaOH (10 M), participation of $(Boc)_2$-Eflornithine occurred. This product was filtered and freeze dried. (32.6 mg; 85 µmol; 20% yield) This Boc protected eflornithine was immediately used for conjugation to [Fe]DAFC by dissolving 8.2 mg (21.4 µmol) of $(Boc)_2$Eflornithine in 500 µL dry DMF and after addition of 1.1 equiv. (8.9 mg, 23.5 µM) HATU, the mixture was left for 10 min at room temperature to activate the carboxylic acid. Subsequently, 0.8 equiv. of [Fe]DAFC (14.2 mg, 16.4 µmol) were added and the mixture was stirred for further 30 min. The end of the reaction was confirmed by analytical RP-HPLC. After removing the organic solvent, product was purified by preparative RP-HPLC. (Gradient A: $t_R$(+Fe) = 10.8/11.2 min). Cleavage solution (TFA:$H_2O$:Triisopropylsilane=95:2.5:2.5) was added to dry product and followed by stirring for 45 min to remove the Boc-protecting groups. After evaporation of the TFA, the product could be isolated via preparative RP-HPLC (gradient 1: $t_R$ (-Fe)= 21.0/21.7 min; $t_R$ (+Fe)= 20.7/21.1 min) and freeze-drying. Analytical data: [Fe]DAFC-eflornithine 1.27 mg [1.2 µmol; 15% of theoretical yield] MALDI TOF-MS: *m/z* [M+H] = 1028.62 [$C_{43}H_{65}F_2FeN_8O_{15}$; exact mass: 1027.86 (calculated)]

[Ga]DAFC-eflornithine 2.4 mg [2.5 µmol; 30% of theoretical yield] MALDI TOF-MS: *m/z* [M+H] = 1040.62 [$C_{43}H_{65}F_2GaN_8O_{15}$; exact mass: 1040.37 (calculated)]

**Conjugation of Fludioxonil**

For the conjugation of fludioxonil butyric acid, 3.8 mg (11.6 µmol) was dissolved in 500 µL anhydrous DMF and after addition of 2 equiv. (8.8 mg, 23.2 µmol) HATU, the mixture was left for 10 min at room temperature to activate the carboxylic acid. Hereafter, 5.0 mg (5.8 µmol) of [Fe]DAFC dissolved in 500 µL DMF were added. The pH was adjusted to 8-9 with DIPEA and rock for 1 hour at room temperature. The end of the reaction was confirmed by analytical RP-HPLC. Crude product was purified by preparative RP-HPLC (gradient 3, $t_R$= 24.4 min) and freeze dried to give a red coloured solid. Analytical data: [Fe]DAFC fludioxonil butyric acid 3.15 mg [2.6 µmol, 49% of theoretical yield] RP-HPLC gradient B, $t_R$ = 12.5 min; MALDI TOF-MS: m/z [M+H] = 1180.32 [$C_{53}H_{65}F_2FeN_8O_{17}$; exact mass: 1179.96 (calculated)].
For [Ga]DAFC-fludioxonil, iron free DAFC-fludioxonil was labelled with stable gallium by adding a 10-fold molar excess of $GaBr_3$ solution at pH 4.5 and stirred for 10 min at room temperature. [Ga]DAFC-fludioxonil 390 µg [2.6 µmol, 35% of theoretical yield] RP-HPLC gradient B, $t_R$ = 12.3 min; MALDI TOF-MS: m/z [M+H] = 1192.79 [$C_{53}H_{65}F_2GaN_8O_{17}$; exact mass: 1192.36 (calculated)].

**Conjugation of Thiomersal**

For the conjugation, thiomersal 3.04 mg (7.5 µmol) was dissolved in 500 µL dry DMF and after addition of 1.1 equiv. (3.1 mg; 8.2 µmol) HATU, the mixture was left for 10 min at room temperature to activate the carboxylic acid. Subsequently, 5.06 mg (5.8 µmol) of [Fe]DAFC were added and the reaction was stirred for further 30 min. End of the reaction was confirmed by RP-HPLC and immediately purified by preparative RP-HPLC (Gradient 3, $t_R$ = 25,27 min) to give a red coloured solid after lyophilisation. Analytical data: [Fe]DAFC-thiomersal 3.26 mg [2.6 µmol, 46 % of theoretical yield]; RP-HPLC gradient B, $t_R$ = 17.51 min; MALDI-TOF-MS: m/z [M+H] = 1229.85 [$C_{46}H_{63}FeHgN_6O_{15}S$; exact mass: 1229.31 (calculated)];
For [Ga]DAFC-thiomersal, iron free DAFC-thiomersal was labelled with stable gallium by adding a 10-fold molar excess of $GaBr_3$ solution at pH 4.5 and the mixture was stirred for 10 min at room temperature. [Ga]DAFC-thiomersal 620 µg [0.5 µmol, 35 % of theoretical yield]; RP-HPLC gradient B, $t_R$ = 17.18 min; MALDI-TOF-MS: m/z [M+H] = 1242.44 [$C_{46}H_{63}GaHgN_6O_{15}S$; exact mass: 1242.03 (calculated)]

**Conjugation of 5-Fluoroorotic acide**

For the conjugation of 5-Fluoroorotic acid, 3.78 mg (21.7 µmol) was dissolved in 500 µL dry DMF and after addition of 1.1 equiv. (9.0 mg, 23.8 µM) HATU and 10 µL DIPEA, the mixture was left for 10 min at room temperature to activate the carboxylic acid. Subsequently, 5.0 mg of [Fe]DAFC (5.7 µmol) were added and the reaction was stirred for further 30 min. The end of the reaction was confirmed by analytical RP-HPLC. (Gradient A: $t_R$([Fe]DAFC-FOA) = 12.65 min) Hereafter, the product was isolated via preparative RP-HPLC (Gradient 1: ($t_R$(+Fe)= 26.68 min) and freeze-dried as a white powder. Analytical data: [Fe]DAFC-FOA 1.74 mg [1.7 µmol; 30% of theoretical yield]; MALDI TOF-MS: *m/z* [M+H] = 1020.24 [$C_{42}H_{56}FeN_8O_{17}$; exact mass: 1019.30 (calculated)]
[Ga]DAFC-FOA 2.96 mg [2.8 µmol; 31% of theoretical yield]; MALDI TOF-MS: *m/z* [M+H] = 1033,38 [$C_{42}H_{56}GaN_8O_{17}$; exact mass: 1032.30 (calculated)]

**Iron free conjugates:**

For iron removal, half of the conjugate was stirred in 1 mL of 100 mM $Na_2EDTA$ solution for 2 hours to produce the iron free compound, which was subsequently purified by preparative RP-HPLC and freeze-dried to get a white solid powder.

DAFC-eflornithine: Analytical data DAFC-eflornithine 0.48 mg [0.4 µmol, >98% purity] analytical RP-HPLC gradient A, $t_R$ = 11.4/11.5 min; MALDI TOF-MS: m/z $[M+H]^+$ = 975.36 [$C_{43}H_{68}F_2N_8O_{15}$; exact mass: 975.04 (calculated)].

DAFC-fludioxonil: Analytical data DAFC-fludioxonil 1.2 mg [1.1 µmol, 99% purity] analytical RP-HPLC gradient B, $t_R$ = 12.6 min; MALDI TOF-MS: m/z $[M+H]^+$ = 1127.32 [$C_{53}H_{68}F_2N_8O_{17}$; exact mass: 1126.46 (calculated)].

DAFC-thiomersal: Analytical data DAFC-thiomersal 1.1 mg [0.9 µmol, 96% purity] analytical RP-HPLC gradient A, $t_R$ = 17.8 min; MALDI TOF-MS: m/z $[M+H]^+$ = 1178.68 [$C_{46}H_{66}HgN_6O_{15}S$; exact mass: 1177.40 (calculated)].

DAFC-FOA: Analytical data DAFC-FOA 0.46 mg [0.47 µmol, 98% purity] analytical RP-HPLC gradient A, $t_R$ = 13.06 min; MALDI TOF-MS: m/z $[M+H]^+$ = 967.24 [$C_{42}H_{59}FN_8O_{17}$; exact mass: 966.39 (calculated)].

#### Antifungal susceptibility tests

**Table S1.** Minimal inhibitory concentration (MIC), defined as no visible growth by the naked eye, of different siderophore conjugates at 24 and 48 h. Results are shown in [µg/mL] as well as [µM] for better comparison. Experiments were conducted in iron-sufficient and iron–depleted *Aspergillus* minimal medium (AMM). (Results reflects the average of three biological replicates. n =3)

| Compound | MIC AMM **(-)Fe** [µg/mL] | | MIC AMM **(+)Fe** [µg/mL] | | MIC AMM **(-)Fe** [µM] | | MIC AMM **(+)Fe** [µM] | |
|---|---|---|---|---|---|---|---|---|
| | *24 h* | *48 h* | *24 h* | *48 h* | *24 h* | *48 h* | *24 h* | *48 h* |
| [Fe]DAFC-eflornithine | >256 | >256 | >256 | >256 | >249 | >249 | >249 | >249 |
| [Ga]DAFC-eflornithine | >256 | >256 | 256 | >256 | >249 | >246 | 246 | >246 |
| Eflornithine | >256 | >256 | >256 | >256 | >1405 | >1405 | >1405 | >1405 |
| [Fe]DAFC-fludioxonil | 128 | 256 | 128 | >256 | 108 | 217 | 108 | >217 |
| [Ga]DAFC-fludioxonil | 16 | 64 | >128 | >128 | 13 | 54 | >107 | >107 |
| Fludioxonil butyric acid | 32 | 64 | 16 | 32 | 96 | 191 | 48 | 2 |
| Fludioxonil | 0.5 | 0.5 | 0.5 | 0.5 | 2 | 2 | 2 | 96 |
| [Fe]DAFC-thiomersal | 0.125 | 0.25 | 0.25 | 0.25 | 0.1 | 0.2 | 0.2 | 0.2 |
| [Ga]DAFC-thiomersal | 0.5 | 1 | 0.5 | 1 | 0.4 | 0.8 | 0.4 | 0.8 |
| Thiomersal | 0.06 | 0.06 | 0.06 | 0.06 | 0.15 | 0.15 | 0.15 | 0.15 |
| [Fe]DAFC-FOA | 256 | >256 | 256 | >256 | 251 | >251 | 251 | >251 |
| [Ga]DAFC-FOA | 16 | 64 | >128 | >128 | 15 | 62 | >124 | >124 |
| FOA | 64 | 256 | 64 | >254 | 368 | 1471 | 368 | >1471 |
| [Fe]DAFC-Cy5 | 256 | >256 | 256 | >256 | 192 | >192 | 192 | >192 |
| [Ga]DAFC-Cy5 | 16 | 32 | 256 | >256 | 12 | 24 | 191 | >191 |
| Cy5 carboxylic acid | 128 | 256 | 256 | 256 | 265 | 529 | 529 | 529 |
| [Fe]TAFC | >256 | >256 | >256 | >256 | >283 | >283 | >283 | >283 |
| [Ga]TAFC | 8 | >256 | >256 | >256 | 9 | >278 | >278 | >278 |

#### Microscopy of antifungal susceptibility assays

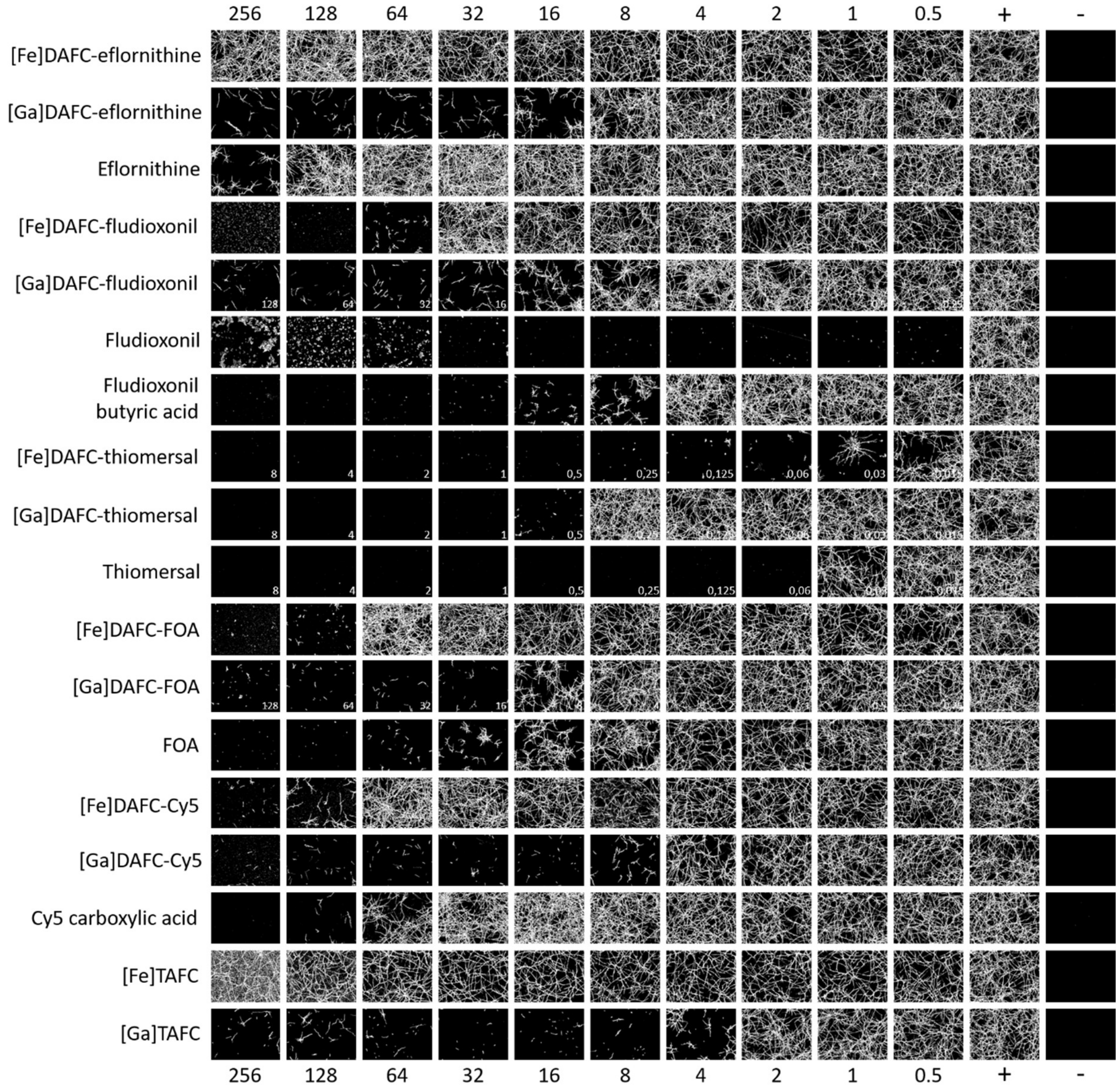


**Figure S3.** Microscopic images of *A. fumigatus* MIC assays with antifungal siderophore conjugates. Number on top and bottom display concentration of the antifungal in [µg/mL]. Images were captured at 24 h incubated at 37°C in **iron depleted** *Aspergillus* minimal medium (AMM Fe(-)). Positive (+) (spores but no antifungal) and negative (-) (just medium) controls are shown in the two columns on the left. Artefacts at 64-256 µg/mL wells of fludioxonil are unsolved crystals of the antifungal.

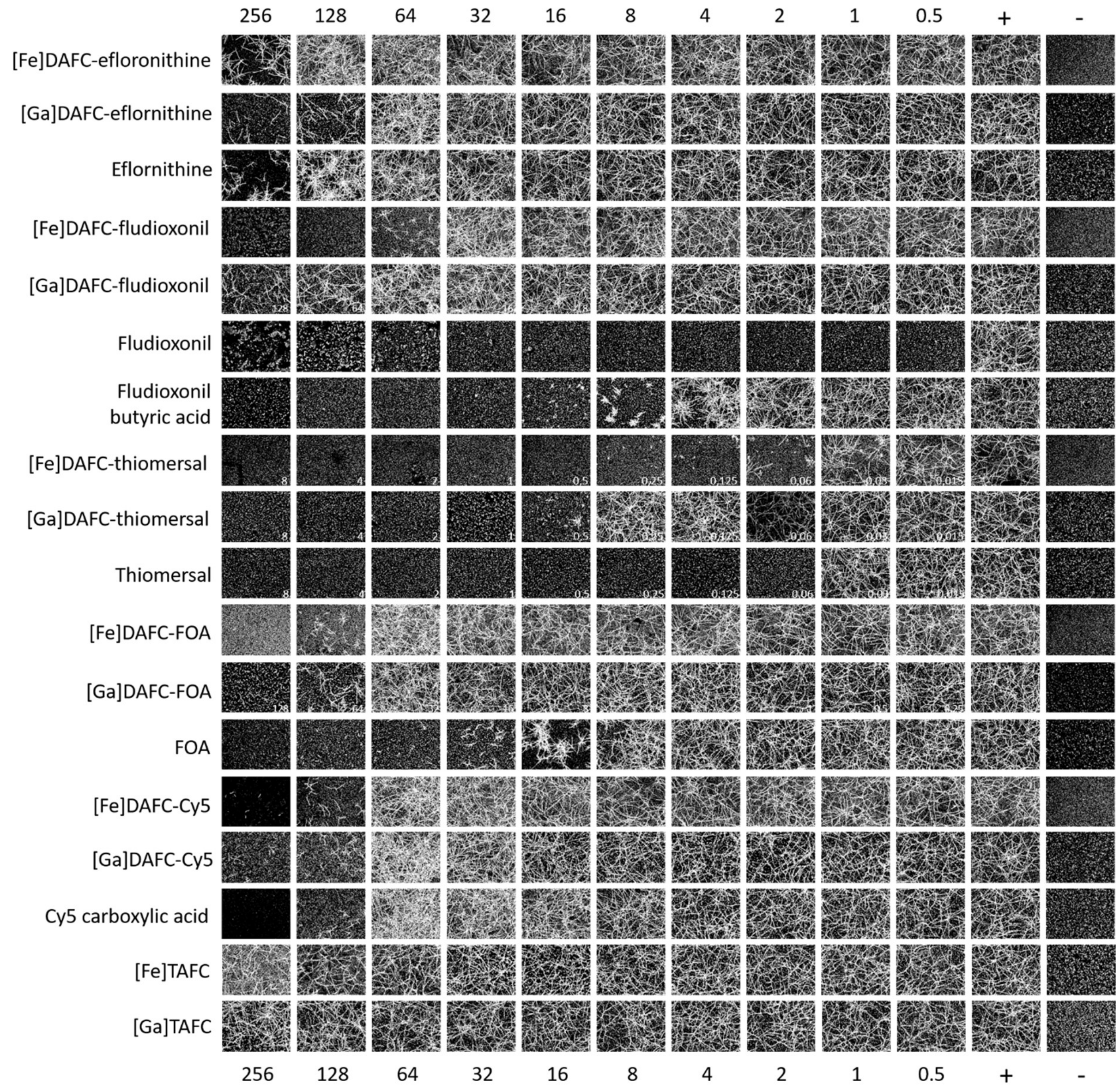


**Figure S4**. Microscopic images of *A. fumigatus* MIC assays with antifungal siderophore conjugates. Number on top and bottom display concentration of the antifungal in [µg/mL]. Images were captured at 24 h incubated at 37°C in **iron sufficient** *Aspergillus* minimal medium (AMM Fe(+)). Positive (+) (spores but no antifungal) and negative (-) (just medium) controls are shown in the two columns on the left. Diffuse punctual background signal origins from the AMM Fe(+) medium.
